# Intrinsic catalytic properties of histone H3 lysine-9 methyltransferases preserve monomethylation levels under low *S-*adenosylmethionine

**DOI:** 10.1101/2022.09.09.507378

**Authors:** Spencer A. Haws, Lillian J. Miller, Diego Rojas La Luz, Vyacheslav I. Kuznetsov, Raymond C. Trievel, Gheorghe Craciun, John M. Denu

**Affiliations:** Wisconsin Institute for Discovery, University of Wisconsin-Madison, Madison, Wisconsin, USA; Department of Biomolecular Chemistry, SMPH, University of Wisconsin-Madison, Madison, Wisconsin, USA; Department of Mathematics, University of Wisconsin-Madison, Madison, Wisconsin, USA; Department of Biological Chemistry, University of Michigan, Ann Arbor, Michigan, USA

**Keywords:** histone, methyltransferase, kinetics, nucleosome, *S*-adenosylmethionine, metabolism

## Abstract

*S*-adenosylmethionine (SAM) is the methyl donor for site-specific methylation reactions on histone proteins, imparting key epigenetic information. During SAM-depleted conditions that can arise from dietary methionine restriction, lysine di- and tri-methylation are reduced while sites such as Histone-3 lysine-9 (H3K9) are actively maintained, allowing cells to restore higher-state methylation upon metabolic recovery. Here, we investigated if the intrinsic catalytic properties of H3K9 histone methyltransferases (HMTs) contribute to this epigenetic persistence. We employed systematic kinetic analyses and substrate binding assays using four recombinant H3K9 HMTs (i.e., EHMT1, EHMT2, SUV39H1, and SUV39H2). At both high and low (sub-saturating) [SAM], all HMTs displayed the highest catalytic efficiency (*k*_cat_/K_M_) for monomethylation compared to di- and trimethylation on H3 peptide substrates. The favored monomethylation reaction was also reflected in *k*_cat_ values, apart from SUV39H2 which displayed a similar *k*_cat_ regardless of substrate methylation state. Using differentially-methylated nucleosomes as substrates, kinetic analyses of EHMT1 and EHMT2 revealed similar catalytic preferences. Orthogonal binding assays revealed only small differences in substrate affinity across methylation states, suggesting that catalytic steps dictate the monomethylation preferences of EHMT1, EHMT2, and SUV39H1. To link *in vitro* catalytic rates with nuclear methylation dynamics, we built a mathematical model incorporating measured kinetic parameters and a time course of mass spectrometry-based H3K9 methylation measurements following cellular SAM depletion. The model revealed that the intrinsic kinetic constants of the catalytic domains could recapitulate *in vivo* observations. Together, these results suggest catalytic discrimination by H3K9 HMTs maintain nuclear H3K9me1, ensuring epigenetic persistence after metabolic stress.

## Introduction

Histone lysine methylation is an epigenetic post-translational modification (PTM) catalyzed by histone methyltransferases (HMTs) on the ε-amino group of histone lysine residues. Histone lysine residues can be utilized as platforms for three distinct methylation states (i.e., mono-, di-, and trimethylation), that along with their locus and surrounding amino acid sequence, confer distinct biological function [^1^]. For example, H3 Lys-4 monomethylation (H3K4me1) identifies active or primed enhancers, while H3 Lys-27 trimethylation (H3K27me3) at gene promoters is associated with transcriptional repression [^2–7^]. The specific site and degree of histone methylation is largely dependent on the depositing HMT(s), enabling tight regulation over a cell’s histone methylation profile [^8^].

Although there are essential functional differences that distinguish HMTs from one another, all require *S*-adenosylmethionine (SAM) as the methyl-donor cosubstrate. SAM is the enzymatic product of a synthetase reaction between the essential amino acid methionine and ATP, which is the only known mechanism supporting intracellular SAM availability in higher eukaryotes [^9,10^]. This reliance on intracellularly derived SAM (via methionine) to support HMT activity creates an interdependence between metabolism and the epigenome [^11, 12^]. Various studies have highlighted this relationship, illustrating how impaired SAM metabolism negatively impacts histone methylation abundance in both isolated cell and whole organism model systems. Interestingly, all forms of histone methylation are not equally sensitive to decreased SAM availability, with di- and trimethylation generally possessing greater sensitivity than monomethylation [^13–15^]. It is hypothesized the sensitivity or robustness of histone methylation to fluctuations in SAM availability may be explained by the inherent catalytic properties of HMTs [^16, 17^]. Biochemical analyses of HMTs report that an increase in substrate methylation states can lead to a corresponding decrease in enzyme catalytic efficiency [^18–24^], but it is unclear how alterations in SAM levels affect the relative rates of mono-, di-, and trimethylation. Therefore, a detailed comparative analysis is needed to comprehensively determine how metabolic fluctuations in SAM availability impact HMT catalysis on substrates with varying degrees of lysine methylation.

In this study, we compare the inherent catalytic properties of four H3 Lys-9 (H3K9) HMTs by employing detailed biochemical analyses that include steady-state and pre-steady state kinetics, and quantitative binding assays, using histone peptide and recombinant nucleosome substrates covering the entire range of H3K9 methylation states (i.e., unmodified through dimethylated). The H3K9 HMTs assessed in this study represent two distinct groups of enzymes: EHMT1/EHMT2 and SUV39H1/SUV39H2. EHMT1 (aka G9a-like protein (GLP) and KMT1D) and EHMT2 (aka G9a and KMT1C) are the primary nuclear H3K9 mono- and di-methyltransferases, best known for regulating facultative heterochromatin and subsequent gene repression [^25–28^]. SUV39H1 (aka KMT1A) and SUV39H2 (aka KMT1B) are reported to catalyze nuclear H3K9 di- and trimethylation, primarily regulating constitutive heterochromatin formation and maintenance [^29–32^]. These functions are critical for the repression of repetitive and retrotransposable elements as well as in regulating 3-dimensional chromosome architecture within the nucleus [^33, 34^]. By (1) assessing the biochemical properties of these enzymes on peptide and nucleosome substrates in a single study and (2) subsequently using these results to develop a mathematical model of nuclear H3K9 methylation dynamics, these results can explain nuclear H3K9 methylation dynamics under rapid loss of intracellular SAM availability.

## Results

### Monomethylation is the most catalytically efficient H3K9 HMT reaction under SAM limitation

We previously demonstrated the levels of monomethylation at H3K9 and H3K27 are maintained during severe intracellular SAM depletion while di- and trimethylation of most histone lysine sites are dramatically reduced [^13^]. Notably, pharmacological inhibition of H3K9 mono- and di-methyltransferases EHMT1 and EHMT2 blocked the ability of cells to sustain both global and loci-specific PTM levels deemed critical for the preservation of heterochromatin stability during this metabolic stress, especially H3K9me1. However, the mechanism(s) by which monomethylation is favored for active deposition under extremely low SAM levels is unknown. One potential explanation for these results is that low intracellular SAM levels favor monomethylation reaction catalysis over high-order states. To address this hypothesis, we determined how SAM availability (i.e., concentration dependence) influences each form of histone lysine methylation catalysis (i.e., mono-, di-, and trimethylation). This was accomplished by assessing the steady-state kinetic properties of recombinantly purified H3K9 HMT catalytic domains that included the canonical mono- and di-methyltransferases, EHMT1 and EHMT2, as well as the canonical di- and tri-methyltransferases, SUV39H1 and SUV39H2.

First, we measured steady-state catalytic rates at varied SAM concentrations and fixed, saturating concentrations of three different H3 peptides covering the entire range of potential substrate methylation states (H3K9un, H3K9me1, and H3K9me2). We fit the resulting data to the Michaelis-Menten equation to obtain the steady-state parameters, *k*_cat_, *k*_cat_/K_M_, and K_M_. At saturating SAM conditions, a comparison of the *k*_cat_ values indicates substrate turnover generally decreases as H3 peptide methylation state increases for EHMT1, EHMT2, and SUV39H1 (Figure 1A and Table 1). The transition from unmodified to monomethylated substrate yields the greatest reduction in EHMT1 and EHMT2 *k*_cat_ rates (i.e., 12.7 to 2.01 min-1 and 7.62 to 2.15 min^-1^, respectively) while SUV39H1 is most negatively impacted by the transition from mono- to dimethylated substrate (i.e., 0.30 to 0.03 min^-1^) (Figure 1C-E). Interestingly, SUV39H2 substrate turnover rates are largely unaffected by H3 peptide methylation status, although *k*_cat_ values for SUV39H2 are consistently lower than those of EHMT1 and EHMT2 when all enzymes are provided with an unmodified substrate (Figure 1F). Thus, at saturating SAM levels, all four enzymes display maximal turnover for monomethylation of the unmodified peptide.

**Figure 1:**
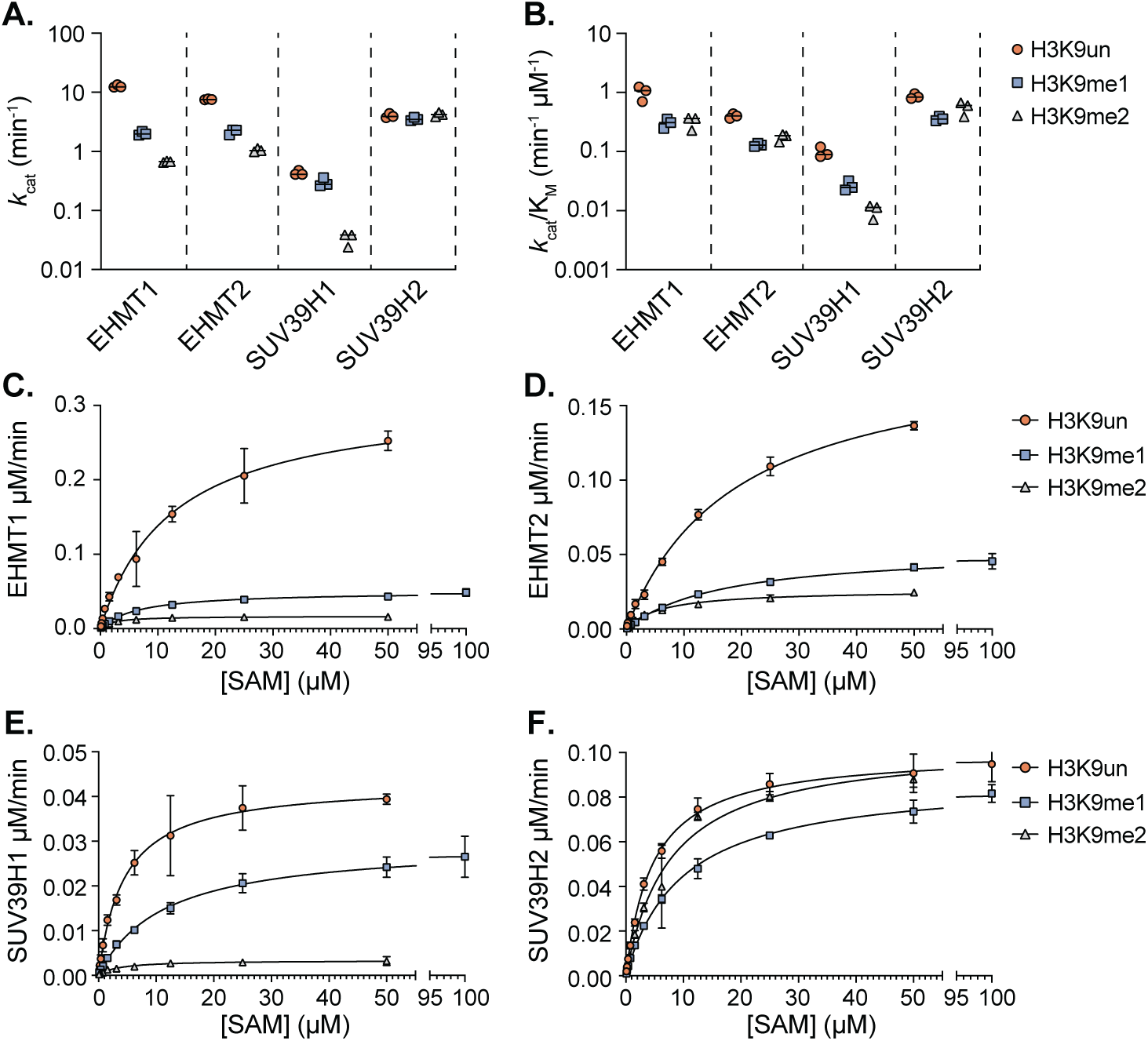
Steady-state kinetic analysis of H3K9 HMTs under varied [SAM] conditions. Summary plots depicting (*A*) *k*_cat_ and (*B*) *k*_cat_/K_M_ values derived from steady-state kinetic analysis of EHMT1, EHMT2, SUV39H1, and SUV39H2 when provided with fixed H3_(1-17)_K9 unmodified (H3K9un), mono-methylated (H3K9me1), or di-methylated (H3K9me2) peptide substrate concentrations and varied SAM concentrations. (*C-F*) Individual steady-state kinetic analyses of EHMT1 (25 nM), EHMT2 (25 nM), SUV39H1 (100 nM), and SUV39H2 (25 nM) under fixed H3_(1-17)_K9 concentrations (50 µM) and varied SAM concentrations (0.1 - 100 µM) which provided the summary values presented in panels (*A*) and (*B*). n=3, error bars represent standard deviation.

**Table 1.**
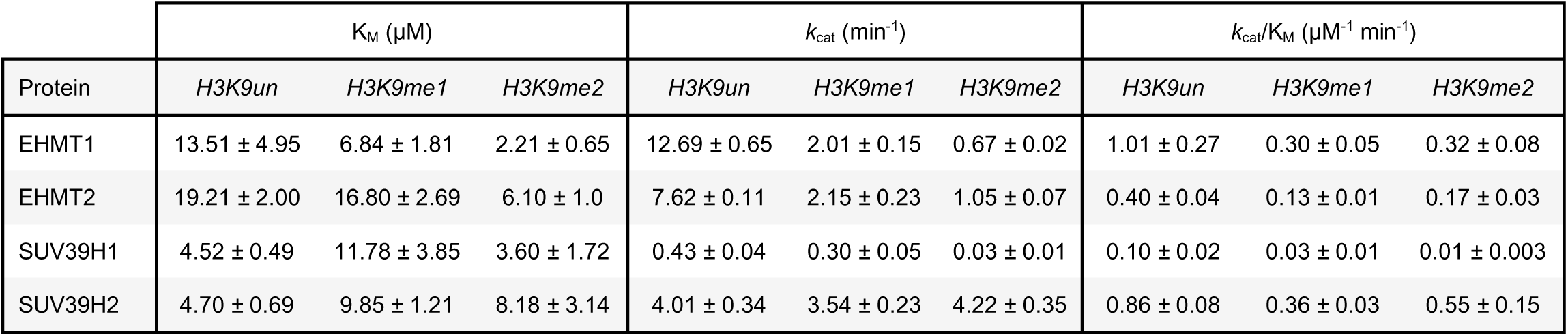
Steady state kinetic constants with SAM as the varied substrate and fixed saturating levels of H3_(1-17)_ K9 peptides.

| Protein | $K_{\text{M}}$ ($\mu\text{M}$) | | | $k_{\text{cat}}$ ($\text{min}^{-1}$) | | | $k_{\text{cat}}/K_{\text{M}}$ ($\mu\text{M}^{-1} \text{min}^{-1}$) | | |
| --- | --- | --- | --- | --- | --- | --- | --- | --- | --- |
|  | *H3K9un* | *H3K9me1* | *H3K9me2* | *H3K9un* | *H3K9me1* | *H3K9me2* | *H3K9un* | *H3K9me1* | *H3K9me2* |
| EHMT1 | 13.51 ± 4.95 | 6.84 ± 1.81 | 2.21 ± 0.65 | 12.69 ± 0.65 | 2.01 ± 0.15 | 0.67 ± 0.02 | 1.01 ± 0.27 | 0.30 ± 0.05 | 0.32 ± 0.08 |
| EHMT2 | 19.21 ± 2.00 | 16.80 ± 2.69 | 6.10 ± 1.0 | 7.62 ± 0.11 | 2.15 ± 0.23 | 1.05 ± 0.07 | 0.40 ± 0.04 | 0.13 ± 0.01 | 0.17 ± 0.03 |
| SUV39H1 | 4.52 ± 0.49 | 11.78 ± 3.85 | 3.60 ± 1.72 | 0.43 ± 0.04 | 0.30 ± 0.05 | 0.03 ± 0.01 | 0.10 ± 0.02 | 0.03 ± 0.01 | 0.01 ± 0.003 |
| SUV39H2 | 4.70 ± 0.69 | 9.85 ± 1.21 | 8.18 ± 3.14 | 4.01 ± 0.34 | 3.54 ± 0.23 | 4.22 ± 0.35 | 0.86 ± 0.08 | 0.36 ± 0.03 | 0.55 ± 0.15 |

Although H3K9 monomethylation was the most enzymatically favorable reaction under saturating levels of both SAM and H3 substrates, it is critical to understand the intrinsic methylation rates when SAM is at very low concentrations such as in cellular conditions of SAM depletion. For this evaluation, we compared *k*_cat_/K_M, SAM_ values amongst each HMT and substrate methylation state combination (Figure 1B, Table 1). The *k*_cat_/K_M, SAM_ parameter is the intrinsic catalytic efficiency of the enzyme when SAM levels are limiting. These *k*_cat_/K_M, SAM_ values revealed the transition from unmodified to monomethylated substrate result in the greatest reduction in catalytic efficiency for EHMT1 (i.e., 1.01 to 0.30 min^-1^ µM^-1^), EHMT2 (i.e., 0.40 to 0.13 min^-1^ µM^-1^), and SUV39H2 (i.e., 0.86 to 0.36 min^-1^ µM^-1^) (Figure 1C-D and 1F). SUV39H1 also displayed this consistent ∼3-fold reduction in catalytic efficiency when transitioning from unmodified to monomethylated substrate (i.e., 0.10 to 0.03 min^-1^ µM^-1^) (Figure 1E). However, SUV39H1 distinctly displayed an additional ∼3-fold reduction in catalytic efficiency when transitioning from mono- to dimethylated substrate while all other enzymes displayed either no change (i.e., EHMT1) or a slight increase (i.e., EHMT2 and SUV39H2).

Together, these data show increased catalytic efficiency for H3K9 mono- relative to di- and trimethylation is a conserved feature of H3K9 HMTs under SAM limiting (*k*_cat_/K_M, SAM_) conditions. Under SAM saturating conditions (*k*_cat_), EHMT1, EHMT2, and SUV39H1 favor monomethylation reactions by 6.4-, 3.5-, and 1.4-fold over dimethylation, and by 19-, 7-, and 14-fold over trimethylation, respectively. In stark contrast, SUV39H2 displayed a similar *k*_cat_ value of ∼4 min^-1^ for all three substrates. These data suggest that for all methyltransferases except SUV39H2, there is strong substrate discrimination that occurs after the enzyme has bound to both SAM and histone peptide.

### H3K9 monomethylation is the preferred reaction under physiologic fluctuations in H3 substrate concentrations

Thus far in the analysis, steady-state kinetic properties suggest H3K9 HMTs intrinsically prefer H3K9 mono- over both di- and trimethylation at both limiting and saturating levels of SAM, reflected in *k*_cat_/K_SAM_ and *k*_cat_, respectively. The notable exception is SUV39H2 which displays no difference in turnover (*k*_cat_) amongst the possible histone substrates. While these conclusions hold at saturating levels of the various methylated peptides, it is important to consider the levels of distinct H3K9 methylation states *in vivo* which fluctuate in response to SAM deprivation. Therefore, it is instructive to determine the intrinsic catalytic efficiency for methylation at non-saturating levels of each H3K9 substrate, which is reflected in the *k*_cat_/K_M_ value for histone substrates. To determine these catalytic efficiency values, we varied the concentrations of each differentially methylated H3K9 peptide at a saturating SAM concentration, determined the steady-state rates for all four HMTs, and fit the data to the Michaelis-Menten equation, obtaining the relevant parameters (Table 2). Analysis of the *k*_cat_/K_M_ values for unmodified, mono- and dimethylated H3K9 substrates reveals a generally larger intrinsic preference for unmodified and monomethylated substrates than observed with *k*_cat_ values among the four enzymes (Figure 2A-B, Table 2). Consistent with the trends in *k*_cat_ values under saturating SAM and H3 peptide substrate conditions, EHMT1 is the most selective for lower methylated substrates while SUV39H2 is the least selective. Together, these data suggest H3K9 HMTs possess an inherent preference for monomethylation when H3 peptide substrate availability becomes limited.

**Figure 2:**
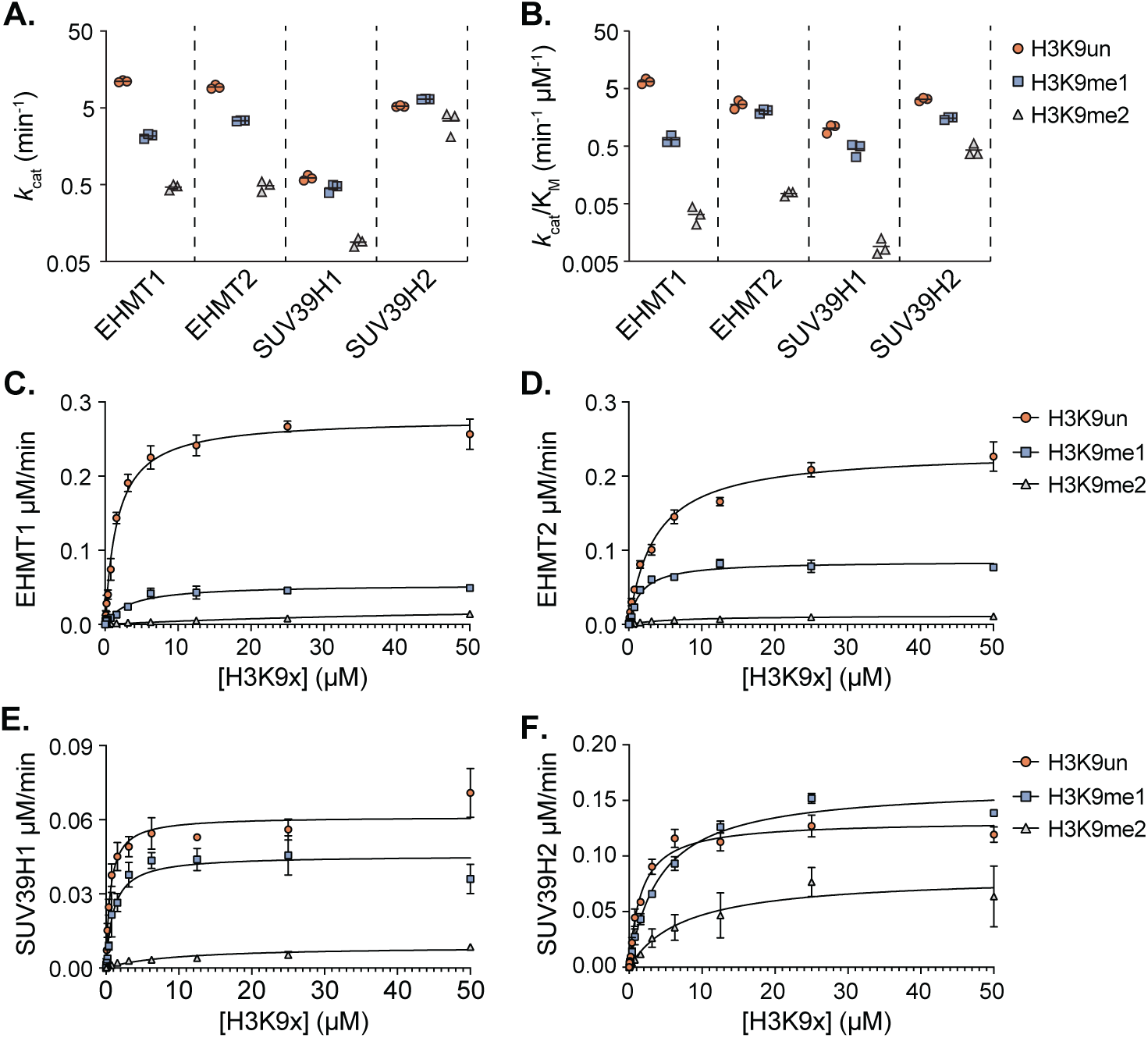
Steady-state kinetic analysis of H3K9 HMTs under H3 peptide substrate limiting conditions. Summary plots depicting (*A*) *k*_cat_ and (*B*) *k*_cat_/K_M_ values derived from steady-state kinetic analysis of EHMT1, EHMT2, SUV39H1, and SUV39H2 when provided with fixed SAM concentrations and varied H3_(1-17)_K9 unmodified (H3K9un), mono-methylated (H3K9me1), or di-methylated (H3K9me2) peptide substrate concentrations. (*C-F*) Individual steady-state kinetic analyses of EHMT1 (25 nM), EHMT2 (25 nM), SUV39H1 (100 nM), and SUV39H2 (25 nM) under fixed SAM concentrations (50 µM) and varied H3_(1-17)_K9 peptide concentrations (0.05 - 50 µM) which provided the summary values presented in panels (*A*) and (*B*). n=3, error bars represent standard deviation.

**Table 2:**
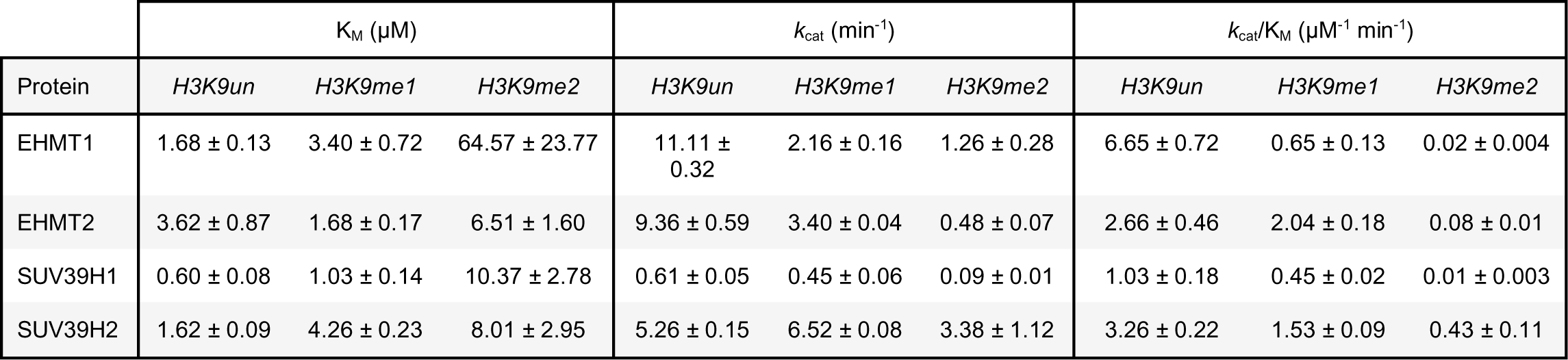
. Steady state kinetic constants with H3_(1-17)_ K9 peptides as the varied substrate and fixed saturating levels of SAM.

To obtain a more physiological perspective on the kinetic preferences for histone methylation targets *in vivo*, we calculated the estimated concentrations of various H3K9 methylation states under SAM abundant and depleted conditions utilizing previously quantified LC-MS/MS HCT116 colorectal cancer cell stoichiometry values for unmodified, mono-, di-, and trimethylated H3K9 residues [^13^]. To do so, we first estimated the concentration of nucleosome-incorporated H3 proteins within the nucleus. This was accomplished by dividing the total base pair (bp) length of the diploid *H. sapiens* reference genome hg38.p13 (including an X and Y chromosome) by 166 bp (146 bp bound + 20 bp linker DNA per nucleosome) to estimate the number of nucleosomes present in a single nucleus. Upon converting the total number of nucleosomes to a mass value of total H3 proteins, we used an estimated nuclear volume for HCT116 cells calculated with a previously published diameter to determine the concentration of H3 proteins within the nucleus [^35^]. Our estimated nuclear H3 protein concentration of ∼170 µM was comparable to those previously reported from other cell types [^36, 37^]. By applying this concentration to our published H3K9 methylation state stoichiometry data, we generated H3K9 methylation state -specific concentrations under both SAM-replete and -deplete conditions (Table 3). This analysis revealed that SAM depletion leads to low- to mid-micromolar fluctuations in H3K9un (i.e., 19.5 to 39.6 µM) and H3K9me2 (i.e., 63.7 to 26.8 µM) abundance while H3K9me1 concentrations remained relatively constant (i.e., 24.7 to 21.1 µM) (Figure S1A-C).

**Table 3:** Estimated nuclear concentrations of H3K9 methylation states.

| | Nuclear Concentration ($\mu\text{M}$) | | | |
| --- | --- | --- | --- | --- |
|  | H3K9un | H3K9me1 | H3K9me2 | H3K9me3 |
| + SAM | $19.5 \pm 3.39$ | $24.7 \pm 4.86$ | $63.7 \pm 8.02$ | $37.7 \pm 4.4$ |
| - SAM | $39.6 \pm 1.36$ | $21.1 \pm 1.53$ | $26.8 \pm 4.96$ | $25.8 \pm 5.7$ |

With good estimates in hand for the full range of methylated H3K9 substrates present under extreme SAM fluctuations, we next looked to determine whether these estimated changes in H3 substrate might impact HMT catalytic rates. HMT catalysis would be reduced if the concentration of H3 substrate dropped below the K_M, H3K9_ values following SAM depletion. We found the K_M, H3K9un_ and K_M, H3K9me1_ values for each enzyme are all at least ∼5-fold below estimated *in vivo* methylated H3K9 concentrations both pre- and post-SAM depletion (Figure 2C-F and Figure S1A-B). The K_M, H3K9me2_ values for EHMT2, SUV39H1, and SUV39H2 are all ≥6-fold below pre-SAM depletion *in vivo* H3K9me3 concentration estimates but never fall more than ∼2.5-fold below our post-SAM depletion estimate (Figure 2D-F and Figure S1C). These observations suggest histone substrate concentrations of all three methylated species are near saturating even under SAM depleted conditions, and importantly, predicts that *in vivo* rates of methylation are dictated by each enzyme’s intrinsic turnover rate when SAM is saturating. Interestingly, the EHMT1 K_M, H3K9me2_ value (i.e., 64.57 µM) is significantly greater than those for each other enzyme (Table 2). This K_M, H3K9me2_ value is similar to pre-SAM depletion H3K9me2 levels (i.e., 63.7 µM) while being greater than post-SAM depletion levels (i.e., 23.77 µM), suggesting EHMT1 is not saturated by this substrate *in vivo*. Therefore, these data suggest low intracellular SAM conditions further disfavor H3K9 trimethylation by EHMT1 although the enzyme is not believed to significantly deposit this form of H3K9 methylation *in vivo*.

### Nucleosomal substrate methylation status influences HMT catalysis similarly to H3 peptides

To assess whether the trends in catalytic efficiency translate to nucleosomal substrates, we performed ‘single turnover’ experiments using recombinant nucleosomes and determined the resulting rate constants. In contrast to steady-state experiments, classical single turnover kinetic experiments use excess enzyme relative to substrate, allowing the enzyme to turnover (i.e., perform catalysis) once. For the subsequent experiments, saturating amounts of enzyme and co-substrate SAM were combined, the reaction was initiated with a low concentration of either unmodified, H3K9me1, or H3K9me2 nucleosomes, and product formation was monitored over time via quantitative immunoblotting (using a standard curve for each probed methylation state). We conducted this experiment with EHMT2 to represent the canonical mono- and di-methyltransferases that would be contributing to the maintenance of H3K9me1 during SAM depletion *in vivo*.

When unmodified nucleosomes were used as the starting substrate, consecutive first-order rate equations were employed to determine the rates (*k*_obs_) of mono-, di-, and trimethylation to account for the ability of EHMT2 to also catalyze di- and trimethylation. In this case, we refer to these methylations as pseudo-single turnover reactions to reflect that multiple sequential methylations occur but under typical single turnover conditions of excess enzyme. For EHMT2, the *k*_obs_ values on unmodified nucleosomes for monomethylation is 2.7-fold greater than dimethylation and 1050-fold greater than trimethylation of H3K9 (0.63 min^-1^, 0.23 min^-1^, and 6E^-6^ min^-1^, respectively) (Figure 3A-B and Table 4). When using H3K9me1 nucleosomes as a starting substrate, the *k*_obs_ values for dimethylation were greater than trimethylation (0.23 min^-1^ and 1.5E^-5^ min^-1^, respectively), consistent with trends and values obtained when using unmodified nucleosome substrates (Figure 3C-D and Table 4). With H3K9me2 nucleosomes as a starting substrate for EHMT2, the *k*_obs_ value for trimethylation was 3.1E^-6^ min^-1^ (Figure 3E-F and Table 4). Consistent with these results, the relative rates of EHMT1 methylation using unmodified nucleosomes displayed similar trends. The EHMT1 *k*_obs_ values on unmodified nucleosomes revealed a modest preference for monomethylation over dimethylation (0.11 (0.13) min^-1^ and 0.096 (0.078) min^-1^, respectively for human and (xenopus) refolded nucleosomes), while there was no significant trimethylation observed over the time course (Figure S2 and Table 4). Therefore, both HMTs, especially EHMT2, exhibit a preference for monomethylation of nucleosomal substrates with a corresponding decrease in *k*_obs_ values as nucleosome substrate methylation status increases.

**Figure 3:**
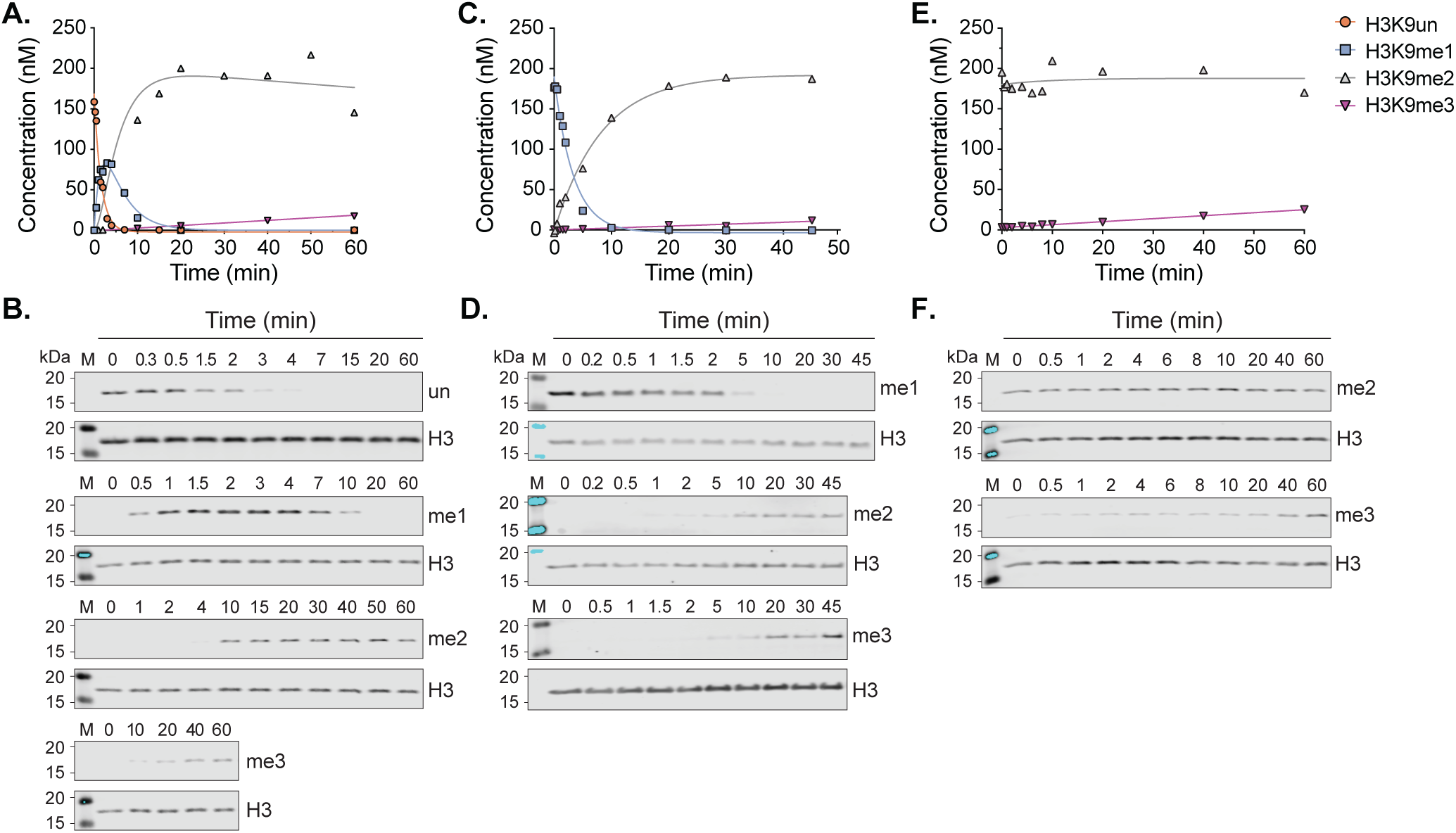
Pre-steady-state kinetic analysis of EHMT2 on nucleosome substrates. Quantification of EHMT2 (40 µM) pre-steady-state experiments with (*A-B*) unmodified, (*C-D*) monomethylated, and (*E-F*) dimethylated recombinant *H. sapiens* nucleosome substrates (100 nM). Representative Western blot images used for the graphical quantifications shown in panels A, C, and E are presented in B, D, and F, respectively. Representative of two experimental replicates. The H3K9 methylation state concentrations were calculated by normalizing the Western blot H3K9 methylation signal by the total H3 signal. Empirically derived concentrations, including those from an additional replicate, were subsequently entered into consecutive reaction first-order rate equations to calculate *k*_obs_ values (See Table 4).

**Table 4:**
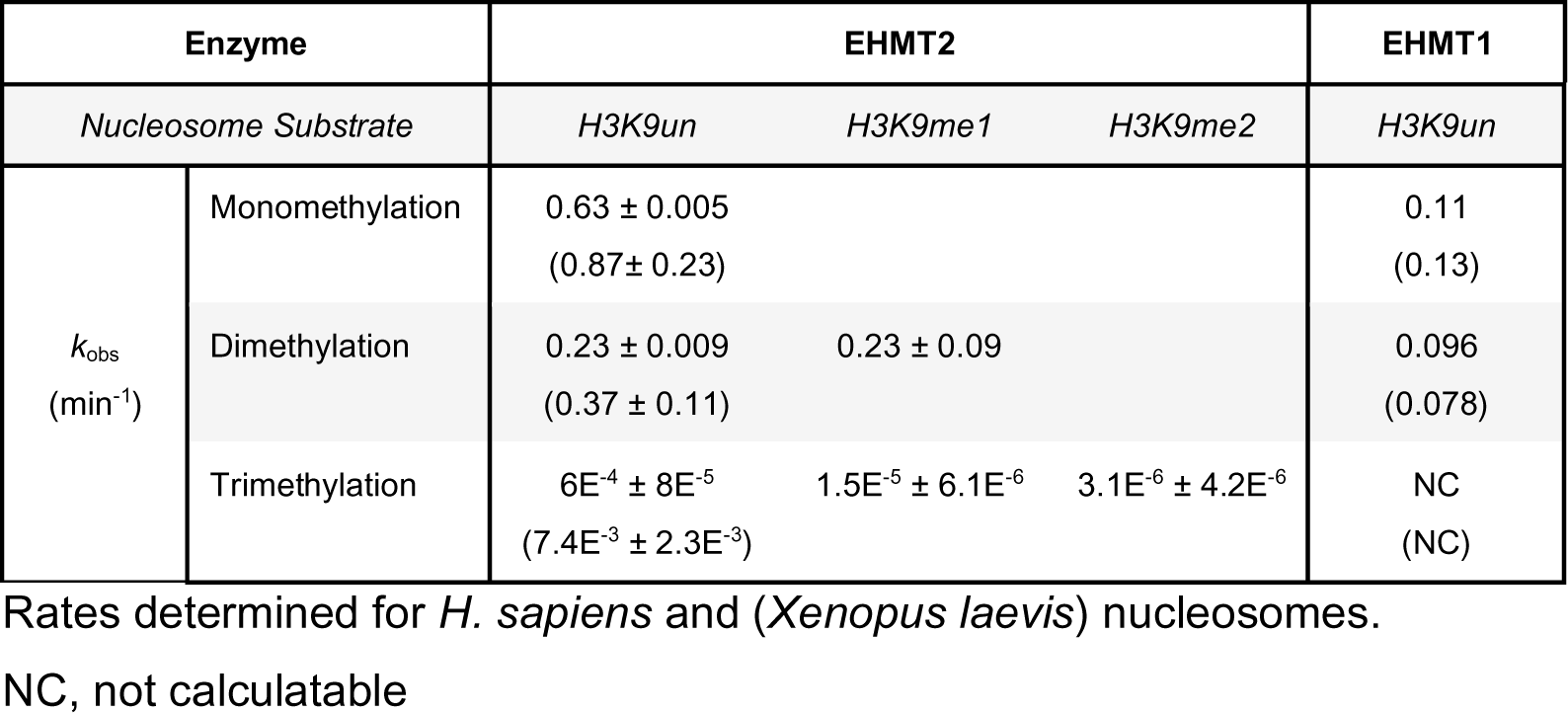
EHMT1 and EHMT2 methylation rates *k*_obs_ for nucleosomes.

Overall, the observed decreases in EHMT1 and EHMT2 *k*_obs_ values as nucleosome substrate methylation status increase demonstrate that nucleosome monomethylation occurs at a faster rate than both the di- and trimethylation reactions. This trend in *k*_obs_ values as a function of nucleosome methylation status is similar to our observed decreases in EHMT1 and EHMT2 *k*_cat_ values when H3K9 peptide substrate methylation status increases. Therefore, these results suggest that the intrinsic catalytic properties of HMT enzymes might explain the observed maintenance of monomethylation when cellular SAM levels dramatically decrease.

### Mathematical modeling guided by HMT kinetic parameters reproduces nuclear H3K9 methylation dynamics during SAM depletion

Given that the intrinsic catalytic properties of HMTs favored monomethylation on both H3 peptide and nucleosome substrates *in vitro*, we next asked whether these empirically-derived kinetic parameters might accurately predict our previously reported *in vivo* H3K9 methylation dynamics [^13^]. To accomplish this, we developed and implemented a mathematical model based on coupled differential equations that describe histone methylation and demethylation when SAM levels drop dramatically.

Our model uses the simplest and most common reaction rate functions, which are given by mass-action kinetics [^38^], as shown below:

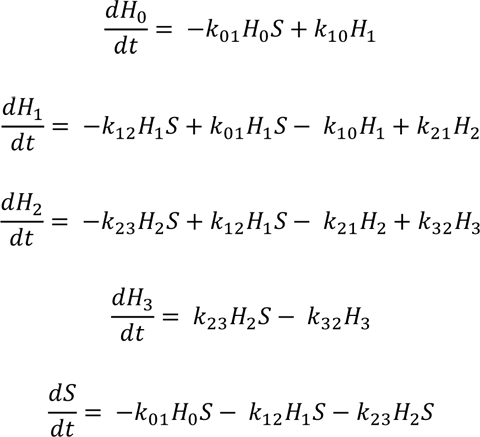

Here, the variables 𝐻_0_, 𝐻_1_, 𝐻_2_, and 𝐻_3_ represent the concentrations of the varying H3K9 methylation states (unmodified, mono-, di-, and trimethylation, respectively), and the variable 𝑆 represents the concentration of SAM. The differential equations take into account that each methylation reaction depends on the concentration of SAM and that one molecule of SAM is consumed at each methylation step. The model parameters (𝑘) are the kinetic parameters (i.e., *k*_cat_/K_M_ or *k*_obs_) for SAM-dependent methylation and demethylation. Importantly, 𝑘_01_, 𝑘_12_, and 𝑘_23_ represent the parameters for mono-, di-, and trimethylation, respectively.

We implemented this mathematical model utilizing the methylation kinetic parameters generated from (1) the steady-state experiments with histone peptides and (2) the pre-steady state experiments with nucleosome substrates to compare to the *in vivo* methylation dynamics data. First, we utilized the *k*_cat_/K_M, SAM_ values for EHMT1 and EHMT2 generated using the steady-state histone peptide experiments to impose a restriction on the ratios of 𝑘_01_ : 𝑘_12_ : 𝑘_23_ to be 3:1:1, which represents a conservative ratio of methylation catalytic efficiencies (Table 1). In parallel, we utilized the *k*_obs_ values obtained from the pre-steady state nucleosome experiments with EHMT2 to impose restrictions on the parameters 𝑘_01_ : 𝑘_12_ to be 3:1, and allowed the last parameter (𝑘_23_) to fluctuate with the model as EHMT2 has an extremely slow trimethylation rate on nucleosomes and may not be representative of cellular trimethylation rates. An additional parameter (not shown in the equations above) allows us to include the possibility that the cell responds to the metabolic stress due to low SAM levels by tapping into a small alternative “emergency supply” of SAM, likely through a methionine salvage pathway. This external source was added as an inflow of a low but constant rate.

This mathematical model generates a numerical simulation that fits experimentally measured histone methylation and SAM levels over a 24-hour time course using both sets of kinetic parameters (i.e., from the steady-state histone peptide and pre-steady-state nucleosome experiments) with the assumption of a small “emergency supply” of SAM, as described above (See experimental methods for information on the estimates/fits for the demethylation rates used in the model) (Figure 4A-B). When using the experimental measurements at time intervals over only the first 90 minutes, the general trends of the simulation results continue to follow the experimental results (Figure 4C-D). Importantly, removing the “emergency supply” of SAM has a significant negative impact on the overall fit of our model over the entire time course, which is reflected in an increase in error between the simulation and *in vivo* data (Figure 4E-F). Overall, this mathematical model highlights the importance of the intrinsic kinetic preferences of the H3K9 HMTs and an accompanying mechanism of a low “emergency supply” of SAM to regulate nuclear H3K9 methylation under severe SAM limitations.

**Figure 4:**
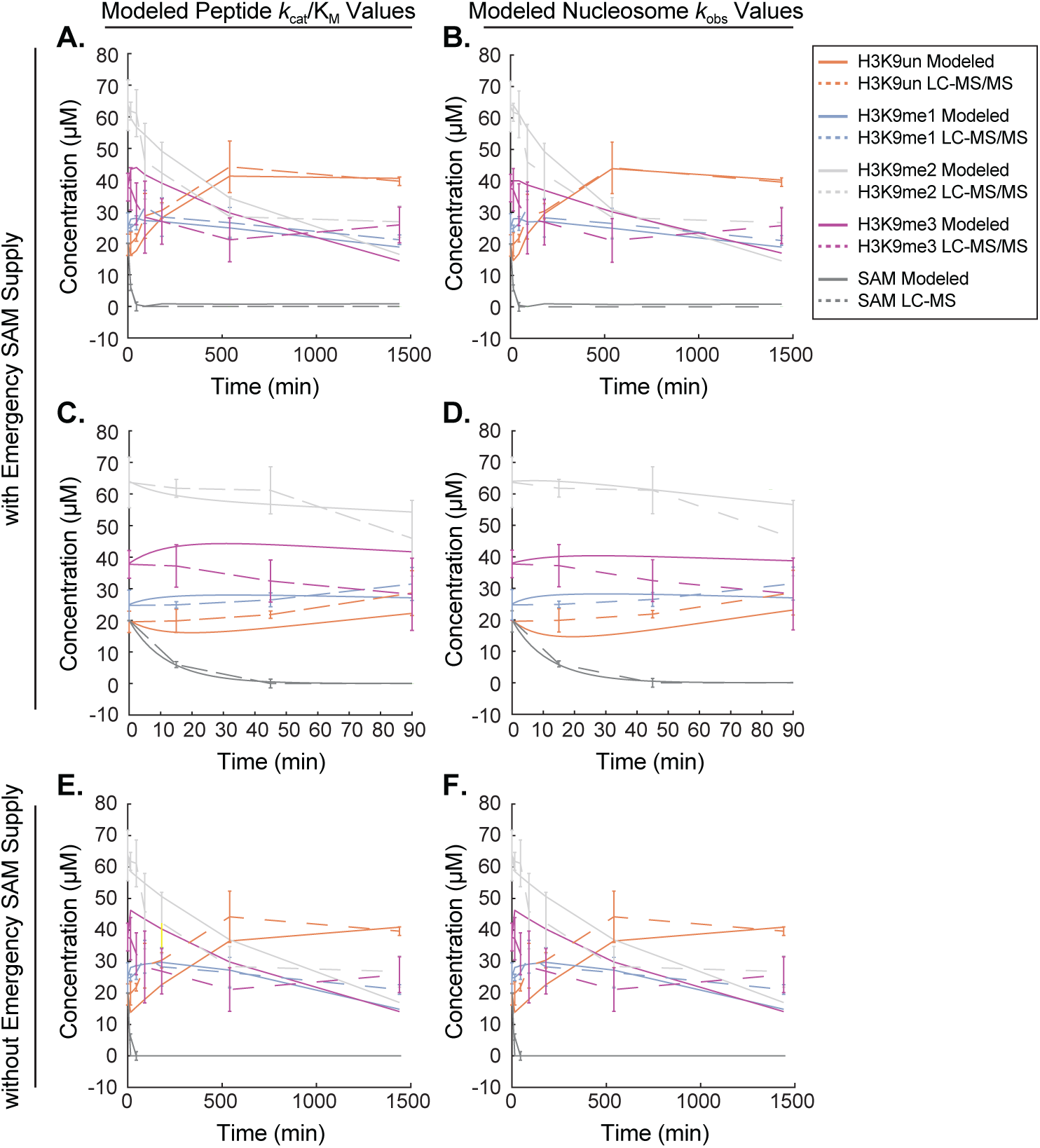
Mathematical modeling of nuclear H3K9 methylation dynamics. Plots depicting changes in H3K9 methylation states and SAM concentrations as measured both empirically via LC-MS-based analyses and by mathematical modeling using the steady-state peptide *k*_cat_/K_M_ data and pre-steady-state EHMT2 *k*_obs_ values generated in this study. A starting SAM concentration of 20 µM was chosen as a representative intracellular value from which relative LC-MS SAM abundance data was converted to µM values. Panels (*A-D*) display modeled H3K9 methylation state and SAM concentrations under the assumption of an existent emergency SAM supply while the modeled data in panels (*E-F*) does not include this assumption. n=3 replicates for empirical H3K9 methylation state and SAM concentration measurements, error bars represent standard deviation.

### Catalytic discrimination underlies HMT preference for H3K9 monomethylation

As H3K9 monomethylation is the most preferred methyltransferase reaction regardless of whether SAM or H3 substrate is limiting, we sought to determine whether substrate methyl-group addition impacts turnover rates through (1) altered binding affinities or (2) via catalytic discrimination. To investigate how methylation of the H3K9 ε-amino group effects binding of the peptide substrate with HMTs, we used fluorescence polarization (FP) assays to generate dissociation constants (K_d_) for each possible combination of differentially methylated H3K9 peptide and HMT assessed in this study. We performed FP assays by titrating enzyme concentrations with fixed concentrations of FAM-labeled H3 peptides in the presence of *S*-adenosylhomocysteine (SAH), the metabolic product of SAM-dependent methyltransferase reactions. Interestingly, we find the methylation state-dependent trends in K_d_ values are distinctly different for EHMT1 and EHMT2 compared to SUV39H1 and SUV39H2 (Table 5). Sequential addition of methyl groups to the H3K9 residue yields a corresponding increase in K_d_ values for both EHMT1 (i.e., H3K9un = 0.029 µM, H3K9me1 = 0.074 µM, H3K9me2 = 0.124 µM) and EHMT2 (i.e., H3K9un = 0.016 µM, H3K9me1 = 0.026 µM, H3K9me2 = 0.04 µM) (Figure S3A-C). However, the relatively small 1.5- to 2.6-fold change increase in K_d_ values following each methyl-group addition are significantly lower in magnitude than the fold change effects on *k*_cat_/K_M_ with increasing peptide methylation of the starting substrate (Figure 1B, Table 1). With the caveat that these binding assays reflect peptide binding to a dead-end product complex with SAH, the data suggest differences in H3 peptide affinity do not account for the larger catalytic efficiency differences for EHMT1 and EHMT2.

**Table 5:** H3_(1-17)_ K9 peptide dissociation constants.

| Protein | $K_d$ ($\mu\text{M}$) | | |
| --- | --- | --- | --- |
|  | *H3K9un* | *H3K9me1* | *H3K9me2* |
| EHMT1 | $0.029 \pm 0.001$ | $0.074 \pm 0.010$ | $0.124 \pm 0.024$ |
| EHMT2 | $0.016 \pm 0.0004$ | $0.026 \pm 0.001$ | $0.04 \pm 0.003$ |
| SUV39H1 | $0.499 \pm 0.045$ | $0.750 \pm 0.047$ | $0.479 \pm 0.123$ |
| SUV39H2 | $1.075 \pm 0.226$ | $0.694 \pm 0.098$ | $0.929 \pm 0.573$ |

Compared to EHMT1 and EHMT2, SUV39H1 and SUV39H2 display almost no effect on H3K9 peptide K_d_ values regardless of methylation status (Table 5). The K_d_ values for both unmodified and dimethylated peptide substrates are comparable (i.e., SUV39H1: H3K9un = 0.499 µM and H3K9me2 = 0.479 µM, SUV39H2: H3K9un = 1.075 µM and H3K9me2 = 0.929 µM) while the monomethylated substrate K_d_ values marginally differ for both enzymes (i.e., SUV39H1: H3K9me1 = 0.75 µM, SUV39H2: H3K9me1 = 0.694 µM) (Figure S3A, D-E). As was the case with EHMT1 and EHMT2, these minor differences in substrate binding affinity upon methyl-group addition are unlikely to impact the *k*_cat_/K_M_ values observed with SUV39H1 and SUV39H2 (Figure 1B, Table 1). Instead, these *k*_cat_/K_M_ and K_d_ values collectively support a model where EHMT1, EHMT2, and SUV39H1 display catalytic discrimination for lower methylation states during chemical catalysis, which holds true at both low and high concentrations of SAM. Therefore, the results presented in this study demonstrate that virtually all intrinsic kinetic parameters of HMTs favor the monomethylation reaction, with monomethylation by SUV39H2 being the unique exception by possessing a catalytic efficiency on par with higher-order methylations.

### Altered active site coordination of the H3K9 ε-amino group suggests catalytic discrimination for differentially methylated substrates

In agreement with our enzymatic comparisons, a structural study of the mutated SET domain lysine methyltransferase SET7/9 Y305F also suggests monomethylation reactions are more catalytically efficient than dimethylation [^23^]. Specifically, Rizzo et al. determined an active-site water molecule positions the unmodified lysine substrate ε-amino group for nucleophilic attack of the SAM methylsulfonium group [^23^]. Displacement of this water molecule prior to dimethylation and subsequent reorientation of the monomethyl lysine substrate have been proposed to underlie the decreased catalytic efficiency of SET7/9 Y305F dimethylation compared to monomethylation of the TAF10 substrate [^23^]. With SET7/9 as a representative model for SET domain catalysis and the Y305F construct mimicking the active site amino acid composition of all HMTs studied here (e.g., EHMT1 F1240, EHMT2 F1152, SUV39H1 F363, and SUV39H2 F370), we asked whether a similar loss of the active site water molecule occurs when EHMT1 is presented with a monomethylated substrate [^22, 39, 40^]. Modeling of the EHMT1 H3K9 monomethylation reaction with existing EHMT1 cosubstrate and product-bound crystal structures revealed the presence of an active site water molecule which, similarly to SET7/9 Y305F, facilitated a network of hydrogen bonds between active site residues and the unmodified peptide substrate (Figure 5A). Specifically, the active site water molecule coordinated hydrogen bonds between active site residues S1196, I1199, and the unmodified lysine ε-amino group, which also shared a hydrogen bond with Y1155. However, when presented with a monomethylated substrate, the active site water molecule was displaced (as seen with SET7/9 Y305F), providing a solvent pocket for the monomethyl lysine to reside, thereby positioning the lysine ε-amino group for another nucleophilic attack of the SAM cosubstrate (Figure 5B). Positioning of the monomethylated lysine ε- amino group was solely mediated through a hydrogen bond with Y1155. These modeling data suggest the greater catalytic efficiency of EHMT1 for H3K9 mono- over dimethylation corresponds with the presence of an active site water molecule-mediated hydrogen bond network that facilitates enhanced coordination of the unmodified H3K9 substrate ε-amino group. Due to the conservation of critical SET domain residues across all HMTs assayed in this study, this mechanism might explain the catalytic discrimination observed. Future studies will be needed to thoroughly investigate the how these HMTs evolved a structure-function relationship that allows the observed catalytic disrimination.

**Figure 5.**
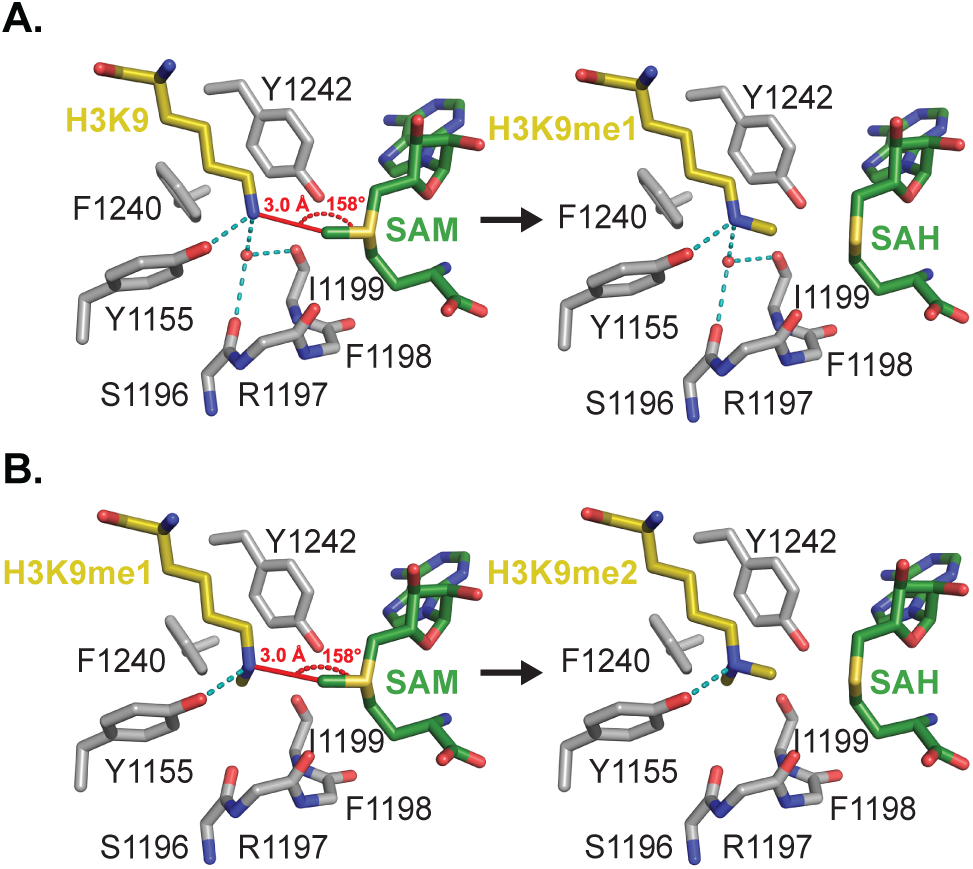
Models for the H3K9 methyl transfer reactions catalyzed EHMT1. (*A*) Model of the monomethyl transfer reaction with H3K9 (yellow carbon atoms) and SAM (green carbon atoms). Key active site residues in EHMT1 and the water molecule that align the K9 ε-amino group for methyl transfer with SAM are depicted with the reaction distance and angle denoted in red. The side chains of residues 1196 – 1199 were omitted for clarity. The right side of the figure illustrates the product complex with H3K9me1 and *S*-adenosylhomocysteine (SAH). (*B*) Dimethyl transfer reaction with H3K9me1 and SAM bound in the active site of EHMT1. Residues, substrates, and the reaction geometry are depicted as in (*A*). The methyl group of K9me1 is bound in the water molecule binding pocket shown in monomethyl reaction complex (*A*), orienting the ε-amino group for the second methyl transfer reaction with SAM. The figure on the right represents the H3K9me2 and SAH product complex.

## Discussion

In a prior study, we found that monomethylation levels of H3K9 were maintained while higher-order methylations dropped during SAM depleted conditions, and concurrent de novo methylation by EHMT1/EHMT2 was required to ensure epigenetic persistence once SAM levels were restored [^13^]. Here, in an effort to understand if the biochemical properties of EHMT1/EHMT2 could explain the *in vivo* dynamics of histone methylation, we performed a detailed biochemical analysis of two functionally distinct pairs of H3K9 HMTs. We found that greater HMT catalytic efficiency for monomethylation relative to di- and trimethylation reactions dictates H3K9 methylation when SAM is limited. In addition, *k*_cat_ and *k*_cat_/K_M_ values for peptides generally favor monomethylation of H3K9. Further, *k*_obs_ values for the canonical mono- and di-methyltransferases on nucleosomes also favor monomethylation over di- and trimethylation of H3K9. These analyses reveal a fundamental property of H3K9 HMTs and suggest an evolutionary pressure to ensure monomethylation of H3K9.

To evaluate the physiological implications of these *in vitro* analyses, we utilized mathematical modeling of nuclear H3K9 methylation state dynamics guided by the kinetic parameters generated here to reproduce our previous *in vivo* observations [^13^]. Together, these data reveal the intrinsic catalytic properties of H3K9 histone methyltransferases are largely sufficient to explain changes in global nuclear H3K9 methylation profiles under severe SAM depletion. Moreover, our mathematical model suggests that the rapid drop in SAM levels following methionine depletion is a direct consequence of H3K9 HMT activity. The model was improved by the inclusion of a small ‘emergency’ source of SAM that could ensure monomethylation for longer periods of methionine restriction. The source of this emergency SAM is unknown, but several possibilities can be envisioned. One source might actually stem from recycling the methyl-groups from protein demethylation. Demethylation reactions produce formaldehyde which can be further metabolized to formate and fed into the folate cycle to promote the regeneration of methionine from homocysteine [^10^]. This methionine can be used as a substrate for SAM synthesis by the systemic mammalian SAM synthesizing enzyme MAT2A [^12^]. Additionally, autophagy-mediated amino acid recycling has also been identified as a highly conserved response to protein restriction from yeast to mammals [^41^]. Such a mechanism could also promote the availability of a small methionine pool for MAT2A-dependent SAM synthesis. Notably, we have previously reported that MAT2A is actively translated in a cell culture model of SAM depletion, providing additional supporting evidence for the existence of an emergency SAM supply when levels of this methyltransferase co-substrate are limited [^13^].

Using both free histone peptides and nucleosomal substrates, we have shown that the H3K9 HMT rate of monomethylation is greater than di- or trimethylation. While these trends hold on both types of substrates, we note that the kinetic parameters determined from the peptide experiments are significantly greater than the *k*_obs_ values determined from the nucleosome experiments. For example, the monomethylation rate for EHMT2 is 10.5-fold greater on peptides than nucleosomes, while the monomethylation rate for EHMT1 is 139-fold greater on peptides. Consistent with our findings, faster rates on peptide substrates compared to nucleosomes have been previously observed. Sanchez et al. reported that an EHMT1/EHMT2 heterodimer possesses a *k*_obs_ ∼800- fold lower for unmodified nucleosomal substrates than for an unmodified free peptide [^21^]. Importantly, the EHMT1/EHMT2 heterodimer utilized by Sanchez et al. possessed similar binding affinities for both mono- methylated peptide (K_d_ = 7.35 µM) and nucleosomal (K_d_ = 2.26 µM) substrates, consistent with our observation that HMT turnover of differentially methylated substrates is driven by inherent catalytic properties toward the histone tail and not through binding preferences [^21^].

One possible explanation for this discrepancy in *k*_cat_ vs *k*_obs_ values for peptide and nucleosomal substrates, respectively, is that turnover of a nucleosome substrate requires a slower additional step during catalysis, before transfer of the methyl group occurs. This prior step in catalysis may be a conformational change in the enzyme and/or nucleosome that facilitates access to the histone H3 N-terminal tail. Previous studies have shown that H3 tail mobility and modifiability within nucleosomes is significantly lower compared to free histone peptides due to the interaction between the H3 N-terminus and nucleosome bound DNA [^42, 43^]. Recently, Morrison et al. reported via NMR that the H3 tails are more conformationally restricted in nucleosomes compared to free histone peptides, binding on either side of the nucleosome DNA dyad and outer DNA turn [^44^]. Therefore, this restriction of the H3 tail within nucleosomes may prompt an extra step to access the H3 tail prior to methylation of H3K9 causing a decrease in the methylation rate on nucleosomes compared to H3 peptides. While this is an appealing explanation, this model would require the conformational shift in tail position to be considerably slower than chemical catalysis, and as such would not be expected to show the same trends in methylation preference as observed here from both free peptides and nucleosomes.

Another plausible explanation is that this slower *k*_obs_ for nucleosome substrates reflects slower chemical rates of methylation compared to peptide substrates. Efficient methyltransferase chemistry involves correct positioning of the two co-substrates for optimal chemical attack of a lone electron pair from the lysine ε-amino group on the SAM methylsulfonium group [^45^]. Furthermore, the attacking lysine ε-amino group must be deprotonated prior to this reaction occurring. It has been proposed that the active site environment and the positive charge on the methylsulfonium of SAM can significantly reduce the lysine p*K*_a_ [^46^]. Therefore, it is possible that the methyltransferase-nucleosome complex affects the active site electrostatics (reduced capacity to lower the ε-amino p*K*_a_) and/or positioning of the H3K9 ε-amino group, leading to less efficient catalysis. Follow up studies will be needed to provide a definitive answer, but regardless, the ordered preference for monomethylation, dimethylation and trimethylation is consistent between peptide and nucleosomal substrates.

To provide a potential explanation for the ability of H3K9 methyltransferases to maintain monomethylation levels under SAM restricted conditions, here we focused on the potential differences between two classes of nuclear HMTs, investigating the catalytic mechanisms of the major *in vivo* contributors to H3K9 mono- and dimethylation (i.e., EHMT1/EHMT2) and di- and trimethylation (i.e., SUV39H1/SUV39H2). Other HMTs not characterized here have also been shown to perform H3K9 monomethylation *in vivo,* such as SETDB1 in addition to the proposed cytoplasmic HMTs PRDM3 and PRDM16 [^32, 47^]. The SET domain of SETDB1 contains all highly conserved residues present in the enzymes assayed in this study, suggesting similar trends in efficient catalysis of H3K9 monomethylation. In contrast, PRDM3 and PRDM16 lack the penultimate tyrosine residue of the consensus ELxF/YDY SET domain sequence which has both structural and catalytic importance in canonical SET domain-containing HMTs [^19, 45, 48, 49^]. Therefore, PRDM3 and PRDM16 require a dedicated study to better understand how their catalytic activity may contribute to meaningful levels of H3K9me1 *in vivo*.

## Experimental procedures

### Recombinant Protein Purification

The *Homo sapiens* EHMT1 catalytic subunit (Addgene plasmid #51314) and *H. sapiens* SUV39H2 catalytic subunit (Addgene plasmid #25115) *Escherichia coli* expression plasmids used in this study were acquired from Addgene, made possible through a gift by Cheryl Arrowsmith. The *H. sapiens* EHMT2 catalytic subunit *E. coli* expression plasmid used in this study was provided by the laboratory of Peter W. Lewis at the University of Wisconsin-Madison. All remaining histone methyltransferase expression plasmids were generated for this study and will be made freely available from Addgene. HMT amino acids included in expression plasmids are as follows: EHMT1 = 982-1266; EHMT2 = 913-1193; SUV39H1 = full length with N-terminal maltose binding protein; SUV39H2 = 162-410.

Transformed Rosetta™ *E. coli* competent cells were cultured in 1 L of 2XYT media at 37°C to an O.D. 600 of 0.8. IPTG was then added to each culture at a final concentration of 1 mM. Cultures were allowed to grow for 16 hours at 18°C before being harvested and stored at −80°C. Pellets were resuspended in 30 mL Buffer A (50 mM NaPi, 250 mM NaCl, 10 mM imidazole, pH 7.8) with protease inhibitors and sonicated on ice with 1 mg/mL lysozyme. Lysate was centrifuged at 45,000xg for 45 min and the supernatant was collected. The supernatant was then loaded onto a HisTrap FF nickel column in line with a GE ÄKTA FPLC. After washing the column with 25% Buffer B (50 mM NaPi, 250 mM NaCl, 250 mM imidazole, pH 7.8), protein was eluted using a linear gradient ending in 100% Buffer B and collected. Fractions were analyzed by Coomassie staining of SDS-PAGE gels to assess purity.

Fractions determined to contain the protein of interest were pooled and dialyzed overnight at 4°C in 4 L of HMT dialysis buffer (20 mM HEPES pH 7.5, 300 mM NaCl, 1 mM TCEP, 10% w/v glycerol). Following dialysis, precipitated protein was pelleted by centrifugation at 4°C for 10 min at 18,000xg and the supernatant was collected. Protein concentrations were determined via absorbance values at 280 nm with extinction coefficient corrections. Final protein samples were aliquoted into single-use 0.2 mL PCR tubes and flash frozen in liquid nitrogen prior to long-term storage at −80°C.

### Recombinant Nucleosome Assembly

Nucleosome-based activity assays were conducted using recombinant nucleosomes made in-house and EpiCypher biotinylated H3K9un-me3 nucleosomes. Recombinant nucleosomes were reconstituted using Widom DNA and *Xenopus laevis* histones with a salt gradient dialysis as established by Luger et al., 1999 [^50^]. Briefly, histones H2A, H2B, H3, and H4 were individually expressed in BL21 (DE3) pLysS competent cells, then purified from inclusion bodies under denaturing conditions by ion exchange chromatography. To refold the histone into octamers, the histones were combined in equimolar ratios and resolved through size exclusion chromatography. Widom DNA was generated from Eco32I (EcoRV) digest from 32-mer inserts in a pUC19 vector. Finally, equimolar histone octamers and DNA were slowly dialyzed from 2 M NaCl to 10 mM NaCl and stored at 4°C.

### Radiometric EZ-Tip Methyltransferase Assay

Michaelis-Menten saturation curves were generated under steady-state conditions with the non-titrated component concentration being present at ≥ 5x K_M_ specific to each enzyme. 1×10^5^ CPM values were provided by [methyl-^3^H] SAM in each reaction. Reaction times and enzyme concentrations were optimized for each HMT to ensure substrate turnover remained below 20%. 50 µM H3 peptide was combined with varying SAM concentrations up to 100 µM and either EHMT1 (25 nM), EHMT2 (25 nM), SUV39H1 (100 nM), or SUV39H2 (25 nM) in histone methyltransferase (HMT) activity buffer (50 mM HEPES pH 7.9, 0.5 mM DTT, 0.1 mM AEBSF, 2 mM MgCl_2_, 0.01% v/v Triton X-100) with a final volume of 20 µL. Enzyme concentrations are calculated for the monomeric form. Reactions were incubated at 30°C for 20 minutes before quenching with TFA for a final concentration of 2% v/v TFA. Then, the reactions were cleaned using StageTips (i.e., two Attract SPE disks (Affinisep) in a pipette tip) with 200 µL acidified water (0.5% v/v formic acid) and eluted with 100 µL elution buffer (80% v/v acetonitrile, 0.5% v/v formic acid). The eluates were added to 2 mL of Ultima Gold™ LSC Cocktail (Sigma-Aldrich) and counted on a scintillation counter (PerkinElmer). All CPM values were corrected for decreased StageTip capacity when peptide concentrations exceeded 10 µM.

### Michaelis-Menten Constant Calculations

CPM values were converted to rates of product formation (µM/min) and used to generate Michaelis-Menten kinetic constants via GraphPad Prism v9.5.0 via the following equation: Y = Et*kcat*X/(Km + X). Variable definitions are as follows: Y = rate of product formation/enzyme velocity; X = substrate concentration; Et = concentration of enzyme in reaction.

### Estimation of In Vivo Histone Concentrations

To estimate the *in vivo* concentrations of H3K9 methylation states, we first estimated the total number of nucleosome-incorporated H3 proteins present within a single nucleus of a *H. sapiens* cell. This was accomplished by calculating the length of the diploid human genome from the haploid GRCh38.p13 assembly. Specifically, we subtracted the lengths of chromosomes X and Y from the total GRCh38.p13 sequence length and multiplied this value by two to generate a diploid length for GRCh38.p13 lacking any sex chromosome inclusion (i.e., 5,772,876,188 bp). We then added back the haploid lengths of chromosomes X and Y to generate a diploid GRCh38.p13 length which includes one of each sex chromosome (i.e., 5,986,144,498 bp). This length was divided by 166 bp, representing the length of nucleosome bound DNA (i.e., 146 bp) as well as a 20bp linker sequence to the neighboring nucleosome, to generate an estimate of 36,061,111 nucleosomes per *H. sapiens* nucleus. Finally, given that two H3 proteins are present within a single nucleus, we simply multiplied our estimated value of nucleosomes per nucleus by two to generate an estimate of 72,122,22 H3 proteins per *H. sapiens* nucleus.

We next began converting our estimate for H3 copies per *H. sapiens* nucleus into a micromolar concentration by first calculating the equivalent mass in grams of H3 proteins per nucleus. To do so, we multiplied our estimate for H3 copies per nucleus by the H3.1 molecular weight of 15,404 Da to calculate a total of 1.1×10^12^ Da of H3 protein per nucleus. As 6.02×10^23^ Da equates to 1 gram of protein, we determined 1.84×10^-12^ g of H3 protein is present within a given *H. sapiens* nucleus. To convert our grams of H3 protein into a concentration, we calculated the volume of an HCT116 nucleus as 6.97×10^-13^ L, using a published HCT116 nuclear diameter of ∼11 µm (Kang et al., 2010) with the required assumption that each nucleus is spherical in shape, and used this value to determine H3 proteins are present at an estimated concentration of 171.85 µM. With our nuclear H3 protein concentration in hand, we simply multiplied this value by our previously published LC-MS/MS H3K9 methylation state stoichiometry values (Haws et al., 2020a) to generate H3K9 methylation state-specific concentrations both pre- and post-SAM depletion in HCT116 cells.

### Nucleosome Pseudo-Single Turnover Assay

Pseudo-single turnover curves with nucleosomes were generated under saturating conditions of enzyme and SAM. 100 nM recombinant nucleosomes was combined with 100 µM SAM (5x K_M, SAM_) and 40 µM enzyme (>800x nucleosome H3 concentration) in histone methyltransferase (HMT) activity buffer (50 mM HEPES pH 7.9, 0.5 mM DTT, 0.1 mM AEBSF, 2 mM MgCl_2_, 0.01% v/v Triton X-100). Enzyme concentrations are calculated for the monomeric form. Reactions were incubated at 30 °C for 60 minutes and quenched with SDS loading dye throughout the time course and then incubated at 95°C for 5 minutes.

The quenched reactions were separated on 15% SDS-PAGE gels, transferred onto 0.2 mm nitrocellulose membranes, and blocked in 50% LI-COR Intercept® (PBS) Blocking Buffer in PBS pH 7.4. Primary antibodies were diluted in 50% LICOR Intercept® (PBS) Blocking Buffer in PBS-Tween pH 7.4 and incubated with the membrane overnight at 4°C. Secondary antibodies were used at a 1:10,000 dilution in 50% LICOR Intercept® (PBS) Blocking Buffer in PBS-Tween pH 7.4 and incubated with the membrane for 1.5 hours at room temperature. Membranes were imaged using an Odyssey Infrared Imager (model no. 9120). Densitometry was performed using Image Studio Lite software.

### Pseudo-Single Turnover Constants Calculations

Densitometry values were converted to concentration of methylation state using a standard curve of H3K9me0-3 nucleosomes and used to generate kinetic constants (*k*_obs_) via GraphPad Prism 9.5.0 via individually and subsequently fitting the following consecutive first-order reaction equations:

For H3K9me0 signal, exponential decay equation:

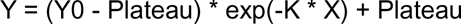

For H3K9me1 signal:

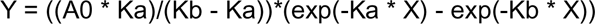

For H3K9me2 signal:

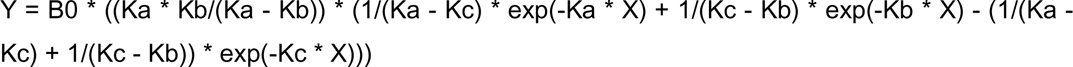

For H3K9me3 signal:

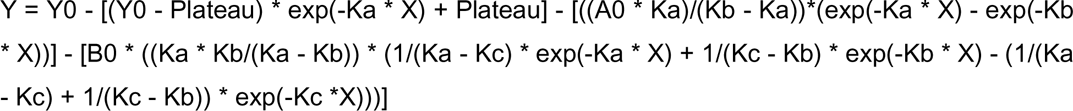

Variable definitions are as follows: Y0 = initial value of unmodified nucleosomes; A0 and B0 = concentration maxima for H3K9me1 and H3K9me2, respectively; X = substrate concentration; Ka = monomethylation rate; Kb = dimethylation rate; Kc = trimethylation rate

### Mathematical Modeling of H3K9 Methylation Dynamics

The MATLAB solver *ode45* was used to calculate numerical solutions to our model which were subsequent inputs for MATLAB’s *lsqnonlin* function together with (1) actual LC-MS/MS measurements (in μM) of H3K9 methylation states as well as (2) estimated LC-MS levels of SAM in HCT116 human cell line following methionine depletion. Both LC-MS/MS H3K9 methylation state stoichiometry data and LC-MS SAM estimates were taken from Haws et al., 2020a [^13^]. Lastly, we used the outputs of the nonlinear least squares optimization algorithm to estimate the values of the parameters in the system, and we used these estimated values to plot the best fit simulation results. The error for the fit is calculated by taking the difference between the numerical simulation solutions and the *in vivo* data, and then dividing by the standard deviation of each time point.

To ensure the validity of the mathematical model, we compared the resulting demethylation rates to those previously determined in the literature ^[50–60]^. In the mathematical model, we fixed the methylation *k*_cat_/K_M_ values to a 3:1:1 ratio for mono:di:trimethylation, reflecting the values obtained for EHMT1, EHMT2, and SUV39H1 with peptide substrates (Figure 4*A*,*C* and Table 1). When using this fixed ratio, the corresponding demethylation *k*_cat_/K_M_ value ratios were modeled to be 8:3:2 for mono:di:trimethylation, respectively. For example, the *k*_cat_/K_M_ value for demethylation of H3K9me3 is twice that of trimethylation. This ratio results in demethylation *k*_cat_/K_M_ values (1.2 ± 1.1, 0.5 ± 0.44, and 0.33 ± 0.30 µM^-1^ min^-1^ for demethylation of H3K9me1, H3K9me2, and H3K9me3, respectively) that are within error of the average values reported in the literature (0.18, 0.16 ± 0.40, and 1.2 ± 2.7 µM^-1^ min^-1^, respectively). Similarly, when we fixed the nucleosome-based *k*_obs_ methylation rates in the mathematical model (0.63 and 0.23 min^-1^ for mono- and dimethylation, respectively, while allowing trimethylation to fluctuate, Table 4), the corresponding demethylation rates were modeled to be 1.4, 0.6, and 0.2 min^-1^ for demethylation of H3K9me1, H3K9me2, and H3K9me3, respectively. These modeled demethylation values are within the first standard deviation of average reported demethylation turnover numbers (*k*_cat_) for demethylation of H3K9me2 and H3K9me3 (2.3 ± 4.9 and 2.4 ± 2.2 min^-1^, respectively). However, while the *k*_cat_ for H3K9me1 demethylation from Goda et al., 2013 is much higher than our modeled rate (16.9 vs. 1.4 min^-1^, respectively), the demethylation turnover number for H3K9me1 still trends as the fastest reaction in agreement with our model.

### Solid Phase Peptide Synthesis

All peptides were manually synthesized on Rink-amide ChemMatrix® resin (Sigma-Aldrich) using DIC/oxyma mediated coupling of 5x molar excess of common Fmoc-protected amino acids for one hour or 2x molar excess of Fmoc-Lys(Boc, Me)-OH (AAPPTec), Fmoc-Lys(Me)_2_-OH-HCl (AAPPTec), Fmoc-Lys(Me)_3_-OH (AAPPTec) amino acids overnight. N-methyl pyrrolidone was used as the primary solvent for all coupling reactions. Completeness of each coupling was confirmed by chloranil test followed by treatment with 10% v/v acetic anhydride in DMF in the presence of 1 M DIEA. Fmoc deprotection was performed with two consecutive 20% v/v piperidine in DMF treatments for 5 min and 15 min, respectively. The N-terminus of all peptides was not capped to improve peptide solubility. All peptides were cleaved with TFA (Reagent K), precipitated with diethyl ether, and C18-HPLC-purified to achieve >99% purity. The molecular weight of each peptide was confirmed using MALDI-TOF mass spectrometry.

### Fluorescence Polarization Peptide Binding Assay

All assay components (i.e., HMT, N-terminus 5-Carboxyfluorescein (5-FAM) labeled peptide, and SAH) were diluted in fluorescence polarization binding buffer (50 mM HEPES pH 7.9, 2 mM MgCl_2_, 0.5 mM DTT, 0.1 mM AEBSF, 0.01% v/v Triton X-100, 10% w/v glycerol). SAH was present at a final concentration of 100 µM while 5-FAM labeled peptides were present at a constant final concentration in the range of 20-100 nM. Monomeric protein concentrations ranged from nanomolar to low-micromolar concentrations (0-10 µM). Reactions were assembled in a black low-flange 384-well plate with a 20 µl final volume. Upon addition of diluted enzyme as final reaction component, 384-well plates were sealed and centrifuged at 1,000xg for 30 sec at room temperature. Prepared 384-well plates were loaded into the BioTek Synergy H4 plate reader after which plates were shaken for 10 sec followed by a 10 min incubation at 25°C. Once the incubation was complete, polarization values were calculated using the BioTek Instruments Gen5 v1.11.5 software package.

### Peptide Dissociation Constant (K_d_) Calculations

Polarization values were exported from BioTek Instruments Gen5 v1.11.5 software and imported into GraphPad Prism v9.5.0 to calculate dissociation constants using the following “One site -- Specific binding” equation: Y = Bmax*X/(Kd + X). Variable definitions are as follows: Y = fluorescence polarization; X = enzyme concentration; Bmax = maximum specific binding; Kd = equilibrium dissociation constant.

### Modeling of EHMT1 Catalyzed Methyl Transfer Reactions

Models for the H3K9 mono- and dimethyl transfer reactions catalyzed by EHMT1 were modeled using the coordinates of the crystal structures of the EHMT1 product complexes bound to SAH and an H3K9me1 peptide (3HNA.pdb) and bound to SAH and an H3K9me2 peptide (2RFI.pdb). SAM was modeled into the active site by superimposing the coordinates of the EHMT1 product complexes with the crystal structure of EHMT1 bound to SAM and a small molecule inhibitor (5TTG.pdb). Figures were rendered and reaction geometries measured using PyMOL (Schrödinger, Inc.).

## Data availability

Full mathematical modeling equations and additional experimental replicates will be shared upon request. Please contact.

## Supporting information

This article contains supporting information.

## Acknowledgements

The authors are grateful to the Peter W. Lewis Lab for providing the EHMT2 plasmid. We are thankful to the members of the Denu Lab for helpful discussions, especially José Moran for assisting with nucleosome generation during assay development.

## Funding and additional information

This work was supported by NIH grant DK125859.

## Abbreviations and nomenclature

EHMT1 /2: [histone H3]-lysine(9) N-trimethyltransferases EHMT1/2, euchromatic histone lysine methyltransferase 1/2
FAM: fluorescein
FP: fluorescence polarization
H3: histone H3
H3K9: histone H3 lysine-9
H3K9me1-3: mono-, di-, and trimethylated H3K9, respectively
H3K9un: unmodified H3K9
HCT116: human colorectal carcinoma cell line
HMT: histone methyltransferase
*k*_cat_: first-order rate constant, turnover number
*k*_cat_/K_M_: catalytic efficiency
K_d_: dissociation constant
*k*_obs_: observed first-order kinetic rate constant
PTM: post-translational modification
SAH: *S*-adenosylhomocysteine
SAM: *S*-adenosylmethionine
SUV39H1/2: [histone H3]-lysine(9) N-trimethyltransferases SUV39H1/2, Suppressor of variegation 3- 9 homolog 1/2

---

**Figure S1:**
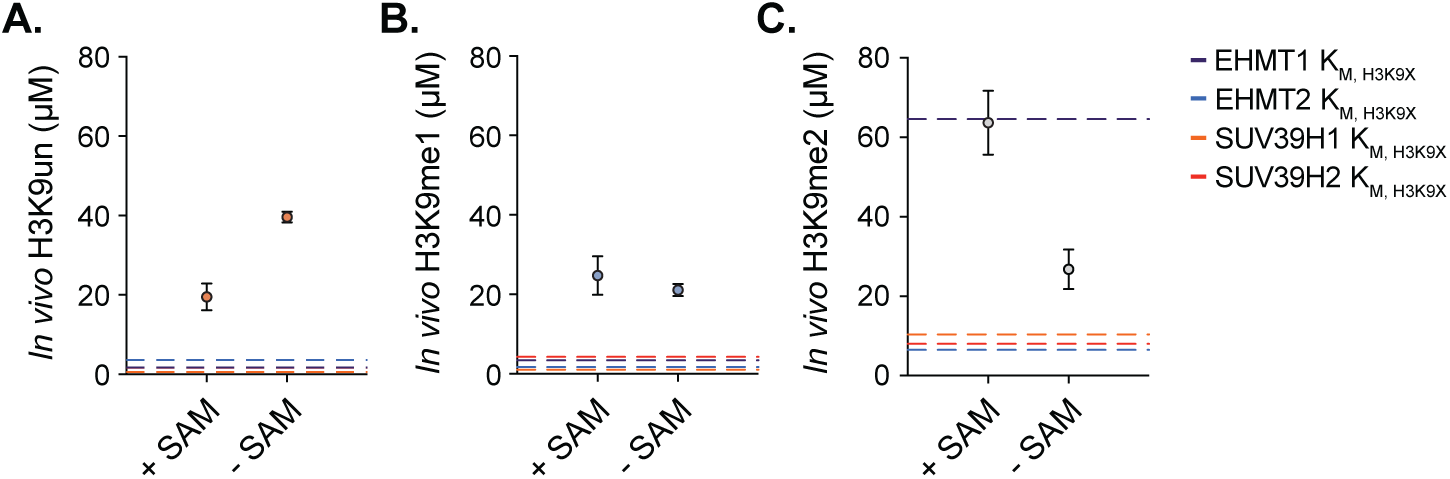
Comparison of HMT K_M, H3K9X_ values with nuclear H3K9 proteoform availability. Nuclear (*A*) H3K9 unmodified (H3K9un), (*B*) monomethylated (H3K9me1), and (*C*) dimethylated (H3K9me2) residue concentrations under SAM-replete and -deplete conditions in HCT116 colorectal cancer cells (See Table 3). H3K9 peptide substrate K_M_ values generated from the kinetic analysis of EHMT1, EHMT2, SUV39H1, and SUV39H2 presented in Figure 3 are superimposed onto each graph to facilitate direct comparison of *in vivo* H3K9 proteoform availability with empirically determined *in vitro* H3K9 peptide proteoform K_M_ values.

**Figure S2:**
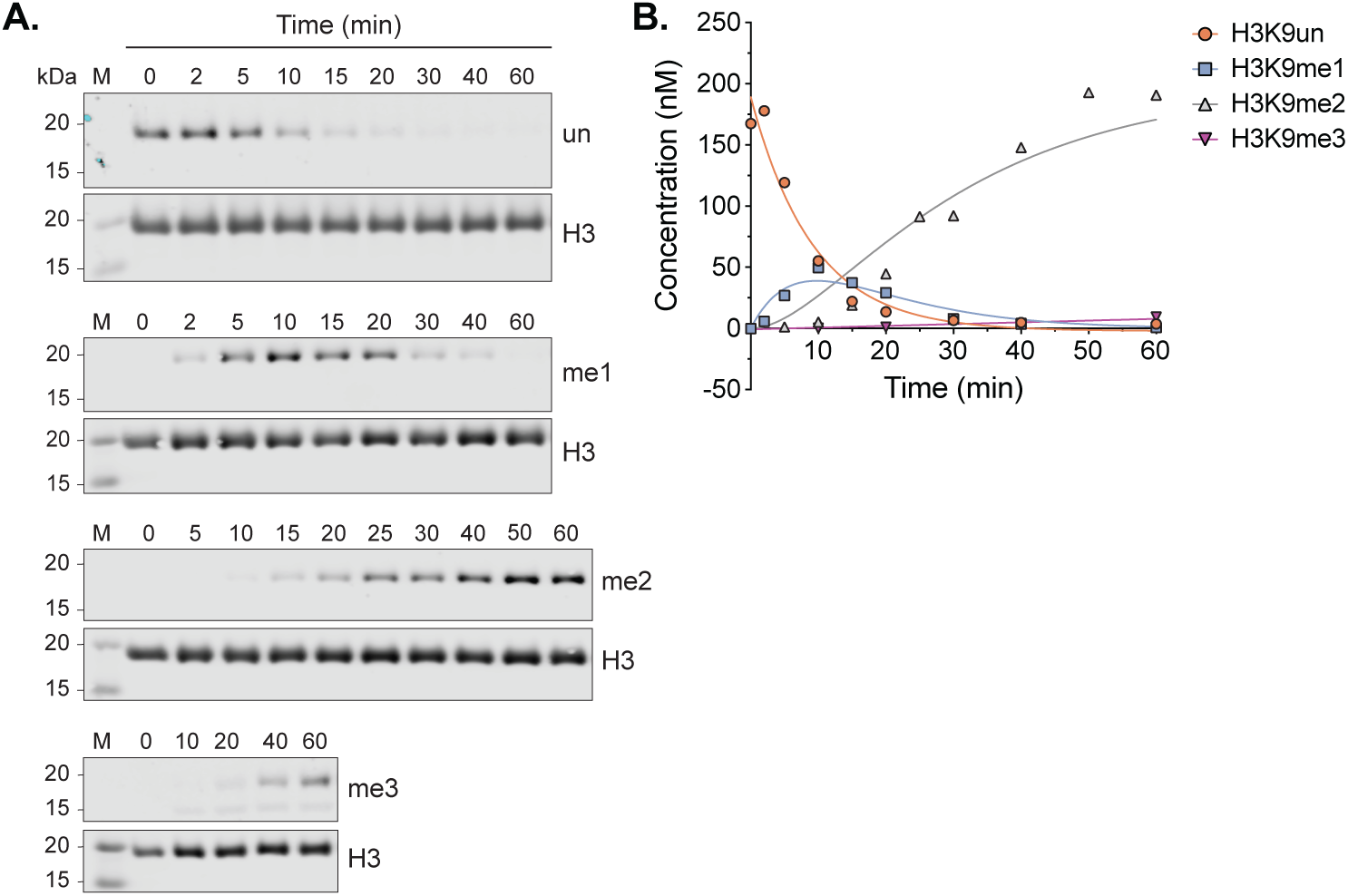
Kinetic analysis of EHMT1 on nucleosome substrates. Quantification of EHMT1 (40 µM) pre-steady-state experiments with (*A-B*) unmodified, recombinant *H. sapiens* nucleosome substrates (100 nM). Western blot images used for the graphical quantifications are shown in panel (*A*). The H3K9 methylation state concentrations were determined by normalizing the methylated K9 signal to the total H3 signal. The *k*_obs_ values were derived from H3K9 methylation concentrations that were fit to consecutive reaction first-order rate equations (See Table 4).

**Figure S3.**
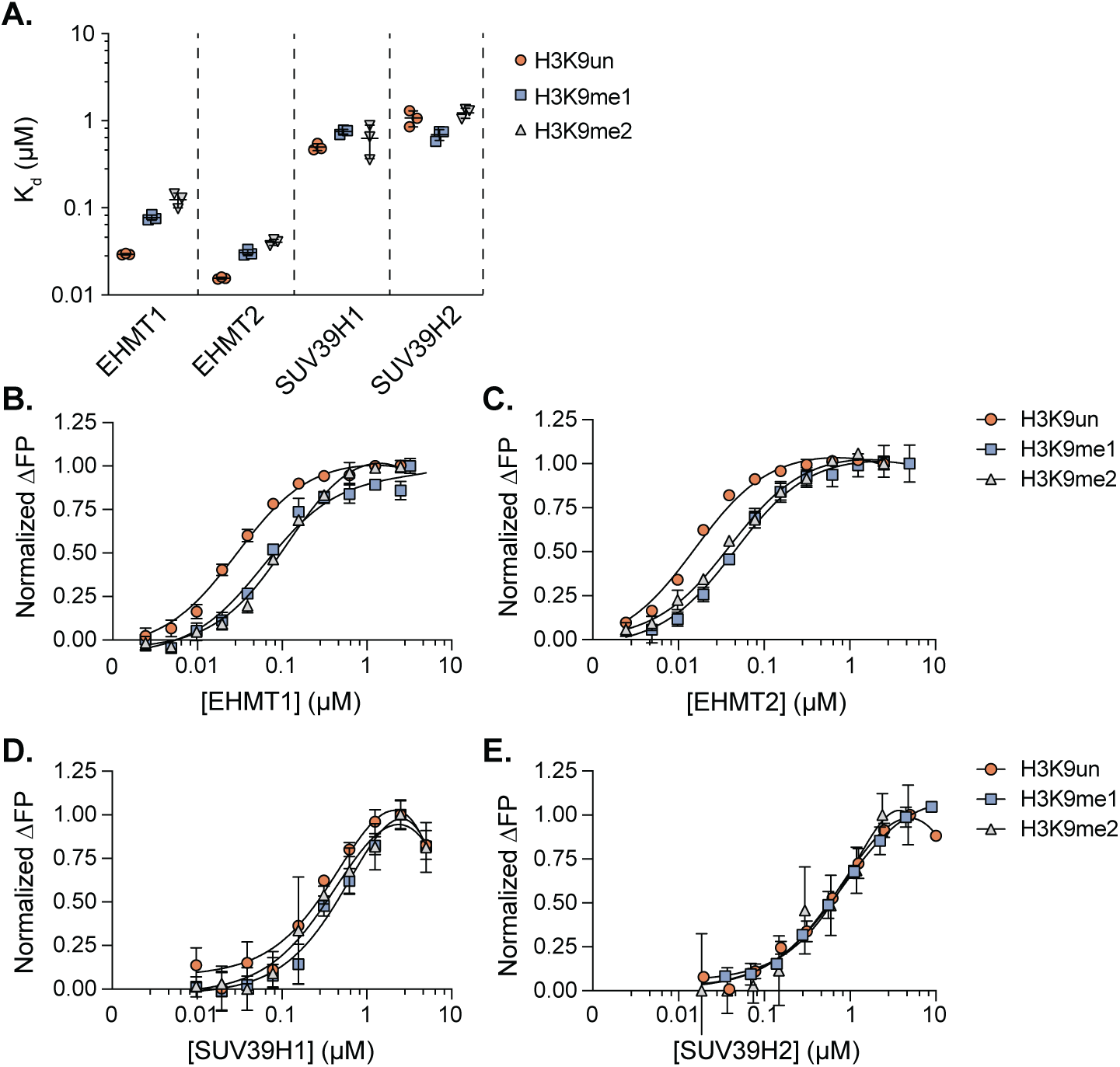
Comparison of HMT H3_(1-17)_K9 proteoform dissociation constants. (*A*) Summary plot depicting K_d_ values derived from Fluorescence Polarization (FP) assays between EHMT1, EHMT2, SUV39H1, and SUV39H2 with 5-FAM labeled H3_(1-17)_K9 unmodified (H3K9un), mono-methylated (H3K9me1), or di-methylated (H3K9me2) peptides. (*C-E*) Individual dose responses of EHMT1, EHMT2, SUV39H1, and SUV39H2 (0-10 µM) under fixed SAH concentrations (100 µM) and fixed H3_(1-17)_ K9 peptide concentrations (varying based on peptide from 20-100 µM) which provided the summary values presented in panels (*A*). Data are presented as average ± s.d. from three technical replicates.

